# Objective Markers of Sustained Attention Fluctuate Independently of Mind-Wandering Reports

**DOI:** 10.1101/2024.07.08.602532

**Authors:** Matthieu Chidharom, Anne Bonnefond, Edward K. Vogel, Monica D. Rosenberg

## Abstract

Sustained attention fluctuates between periods of good and poor attentional performance. Two major methodologies exist to study these fluctuations: an objective approach that identifies "in-the-zone" states of consistent response times (RTs) and "out-of-the-zone" states of erratic RTs and a subjective approach that asks participants whether they are on-task or mind wandering. Although both approaches effectively predict attentional lapses, it remains unclear whether they capture the same or distinct attentional fluctuations. We combined both approaches within a single sustained attention task requiring frequent responses and response inhibition to rare targets to explore their consistency (N=40). Behaviorally, both objective out-of-the-zone and subjective mind-wandering states were associated with more attentional lapses. However, the percentage of time spent out-of-the-zone did not differ between on-task and mind-wandering periods and both objective and subjective states independently predicted error-proneness, suggesting that the two methods do not capture the same type of attention fluctuations. Whereas attentional preparation before correct inhibitions was greater during out-of-the-zone compared to in-the-zone periods, preparation did not differ by subjective state. In contrast, post-error slowing differed by both objective and subjective states, but in opposite directions: slowing was observed when participants were objectively out-of-the-zone or subjectively on-task. Overall, our results provide evidence that objective and subjective approaches capture distinct attention fluctuations during sustained attention tasks. Integrating both objective and subjective measures is crucial for fully understanding the mechanisms underlying our ability to remain focused.

## INTRODUCTION

Sustained attention is the capacity to maintain focus on a task over an extended period. This ability is crucial for our daily activities, yet it proves challenging to keep a consistent level of attention throughout long durations. Attention fluctuates continuously, alternating between phases of high and low engagement. During driving, for example, moments of low attention can manifest objectively in performance outcomes, such as missing a highway exit. They can also be associated subjectively with mind-wandering, or the experience of task-unrelated thoughts (i.e., thinking about the upcoming vacation plans). This scenario illustrates the two methodologies traditionally employed to isolate these fluctuations: one objective, relying on behavioral performance, especially reaction time variability (Esterman et al., 2013), and the other subjective, based on self-reported unrelated thoughts (McVay & Kane, 2009). Although both approaches are effective in identifying variations in sustained attention and isolating error-prone mental states, they have traditionally been studied separately. One might assume that both objective and subjective methodologies reflect the same attentional fluctuations, but is this truly the case?

One objective methodology is a performance-based approach that examines the within-subject variability of response times (RTs) on sustained attention tasks that require frequent responses to non-target stimuli and response switches or inhibitions to rare targets (Esterman et al., 2013; Rosenberg et al., 2013). In this approach, each participants’ response variability is measured with the variance time course (VTC), a metric computed as the absolute deviance of each correct-trials RT from the mean RT on the task. A median split of each individual’s VTC distinguishes periods of low RT variability or consistent responding (‘in-the-zone’ states) and periods of high RT variability or erratic responding (‘out-of-the-zone’ states). Variable out-of-the-zone states are more error-prone compared to consistent in-the-zone states. Such effects have been consistently replicated across different sustained attention tasks (Chidharom, Krieg, Pham, et al., 2021; Esterman et al., 2014, 2015; Esterman, Poole, et al., 2017; Esterman, Thai, et al., 2017) and also in clinical populations (Chidharom, Krieg, & Bonnefond, 2021), making the VTC an efficient and objective manner to isolate attentional states. The second methodology for measuring attention fluctuations is a subjective approach. It isolates fluctuations of sustained attention by employing thought-probes that repeatedly query participants about their thought content during a sustained attention task. Historically, two primary states have been identified: an ‘on-task’ state during which thoughts are task-oriented and an ‘off-task’ state (or mind-wandering states) in which participants report experiencing more task-unrelated thoughts. Research has shown that there is a higher frequency of commission errors during mind-wandering periods compared to on-task periods (Andrillon et al., 2021; Bastian & Sackur, 2013; Kane & McVay, 2012; McVay & Kane, 2009; Seli et al., 2013; Stawarczyk, 2014; Stawarczyk et al., 2013, 2020), suggesting that thought-probe-based approach is a reliable and valid way to isolate attentional states (Kane et al., 2021; Welhaf et al., 2023).

Although these two methods effectively isolate attentional fluctuations, it remains unclear to what extent they measure the same or distinct attentional fluctuations. An intuitive hypothesis is that both methods are consistent and measure the same type of fluctuations given their similar behavioral patterns: periods of self-reported mind wandering and greater RT variability are both associated with increased error rates. Efforts to reconcile these methods, such as those by Bastian & Sackur (2013), have shown that the temporal position of a thought probe relative to the nearest local peak of RT variability can predict mind wandering. Similarly, recent studies using the metronome response task concluded that behavioral variability is an online marker of off-task thoughts and mind wandering (Godwin et al., 2023; Seli et al., 2013). Thus, according to the *hypothesis of consistency*, the objective and subjective methods measure the same type of fluctuation.

However, recent empirical evidence fails to corroborate this viewpoint, thereby lending support to an alternative *hypothesis of inconsistency*. Notably, Unsworth et al. (2021) conducted an inter-subject latent variable analysis which revealed that reaction time variability on the Sustained Attention to Response Task (SART; Robertson et al., 1997), operationalized by the coefficient of variation, and self-reported task-unrelated thoughts represent two independent and distinct measures. These findings suggest that RT variability and mind wandering reflect separate cognitive processes or constructs rather than manifestations of a unitary attentional fluctuation phenomenon. The *hypothesis of inconsistency* is further supported by fMRI evidence of paradoxical underlying neural mechanisms (Kucyi et al., 2016). Both in-the-zone periods of good performance and off-task periods of poor performance show greater DMN activity than out-of-the-zone and on-task periods, respectively (Christoff et al., 2009; Esterman et al., 2013, 2014; Hasenkamp et al., 2012a; Kucyi et al., 2016, 2023; Mittner et al., 2014). Work by Kucyi and colleagues (2016) has tried to distengle the variance underlying DMN activity and revealed that both off-task thoughts (low efficiency state) and behavioral RT stability (high efficiency state) related to DMN activity, suggesting a multidimensional pattern of attentional fluctuations.

The inconsistency hypothesis of subjective and objective approaches could also elucidate the contradictory findings of recent years regarding the role of cognitive control in sustaining attentional performance. Indeed, most theories of sustained attention, such as the resource control theory proposed by Thomson (2015), posit that the gradual reduction of cognitive control over time accounts for declining performance and the automatic redirection of thoughts toward mind-wandering (Thomson et al., 2015). This pivotal role of cognitive control has long been supported by neural data obtained through the subjective approach. Periods of subjectively reported on-task states have been associated with greater engagement of the dorsal attention network (DAN) and the frontoparietal control network (FPCN) compared to off-task periods (Christoff et al., 2009; Hasenkamp et al., 2012b; Kucyi et al., 2016; Mason et al., 2007; Mittner et al., 2014). A similar pattern has been observed from an electroencephalographic (EEG) perspective. Chidharom and colleagues (2023) engaged participants in the SART and explored frontal theta (4-8 Hz) oscillations as a marker of cognitive control engagement (Cavanagh et al., 2009, 2012). The authors found higher frontal theta phase coherence over frontal midline electrodes on trials preceding a critical no-go stimulus, but only during on-task periods and not mind-wandering episodes. This suggests that the subjective fluctuations of the mind correlate with the impaired cognitive control engagement (Chidharom & Bonnefond, 2023).

Although the subjective approach aligns with the current theories of sustained attention, the objective approach instead associates decreased cognitive control with *better* attentional performance. Behaviorally, RTs are slower before correct vs. error rare target trials during out-of-the-zone periods but this effect is mitigated when participants are in-the-zone (Esterman et al., 2013). This suggests that correct inhibition of response to rare target trials relies on enhanced preparatory attention, especially during periods of high RT variability. These behavioral findings are supported by fMRI data showing that, when participants are out-of-the-zone but not in-the-zone, DAN activation is higher and errors are preceded by reductions in DAN activity (Esterman et al., 2013). Better understanding the extent to which RT variability and mind wandering probes measure similar or different attentional fluctuations would allow for better characterization of the role of cognitive control in sustained attentional performance.

While both methods have been used separately so far, the goal of this study is to explore the consistency between objective measures and subjective reports of engaged attention. To do so, we combined objective VTC and subjective mind-wandering probes within a single experiment, using, for the first time, a robust intra-individual design to examine whether these approaches identify the same or different type of attentional fluctuations. Our first approach focuses on the time spent out-of-the-zone. Under the consistency hypothesis, this duration should be significantly longer during off-task than on-task periods, suggesting both methods track the same fluctuations. Conversely, if the inconsistency hypothesis is valid, the duration out-of-the-zone should not differ significantly between off-task and on-task periods. Our second approach characterized the predictors of attentional lapses (i.e., failures to inhibit responses to rare no-go stimuli). According to the consistency hypothesis, self-reported mind wandering and RT variability should predict errors with a high degree of collinearity. In the event of inconsistency, the two measures should predict errors independently. Finally, we tried to better characterize the role of cognitive control through the analysis of RTs before and after no-go trials. We assessed both preparatory control before a correct inhibition and slowing adaptation after an error. We expect to find greater pre-correct slowing in out-of-the-zone compared to in-the-zone periods, consistent with earlier findings. If consistency prevails, greater pre-correct slowing should be observed during off-task periods, whereas if the inconsistency hypothesis prevails, preparatory control should be observed during on-task periods. We applied similar logic to the adaptive post-error slowing mechanism. Here we predicted higher engagement of cognitive control during the combined on-task/out-of-the-zone state because both states have been associated with increased control. This study uses a robust intra-subject design in this context, enabling us to gather evidence to determine whether objective and subjective methods measure congruent aspects of attentional fluctuation.

## METHOD

### 2.1. Participants

The dataset analyzed comprised data from 47 individuals, drawn from open-access resources previously published by Chidharom & Bonnefond (2023) on the subject of mind-wandering. The data are available on the Open Science Framework repository (https://osf.io/xwe23/) and was not pre-registered. After the exclusion of seven participants due to incomplete datasets or absence of mind-wandering reports, the study proceeded with a final sample of 40 participants. The average age within this cohort was 22.47 years (SD = 2.21) and the group included 30 male and 10 female participants. Selection criteria were stringently applied to ensure the exclusion of any individual with a history of neurological disorders, those currently undergoing benzodiazepine treatment, individuals with recent drug abuse, or those who had received general anesthesia in the preceding three months. Written informed consent was secured from each participant before their involvement in the study, which adhered to the ethical guidelines of the Helsinki Declaration. Participants were compensated for their contribution to the research.

### 2.2. Experimental design

Participants engaged in the sustained-attention-to-response task (SART) as described by Robertson et al. (1997). During the SART, a random sequence of single-digit numbers from 1 to 9 was presented, each for a brief duration of 150 milliseconds, with each number having an equal chance of appearing (1/9 odds). The task consisted of 720 go trials and 90 no-go trials. Each number was followed by one of five interstimulus intervals (ISIs), selected at random following a uniform distribution: 1500 ms, 1700 ms, 2100 ms, 2300 ms, or 2500 ms. To increase cognitive load, numbers were shown in five different font sizes of Arial typeface: 100, 120, 140, 160, and 180, designed to prioritize the cognitive processing of numerical value over the identification of any peripheral feature of the no-response target digit. The corresponding vertical visual angles of these font sizes were 1.39 degrees, 1.66 degrees, 1.92 degrees, 2.18 degrees, and 2.45 degrees. Digits were displayed in black against a gray background, positioned 0.25 degrees above a central yellow fixation cross, on a standard 17-inch computer monitor. The digits, including the number 3, were shown with equal frequency from a viewing distance of 70 centimeters. Subjects were asked to react swiftly to all digits presented (go trials), with the exception of the digit 3 (no-go trial), where they were to withhold their response. The total duration of the task was around 30 minutes. Participants completed a brief training block lasting two minutes prior to starting the main task (**Figure 1**).

**Figure 1.**
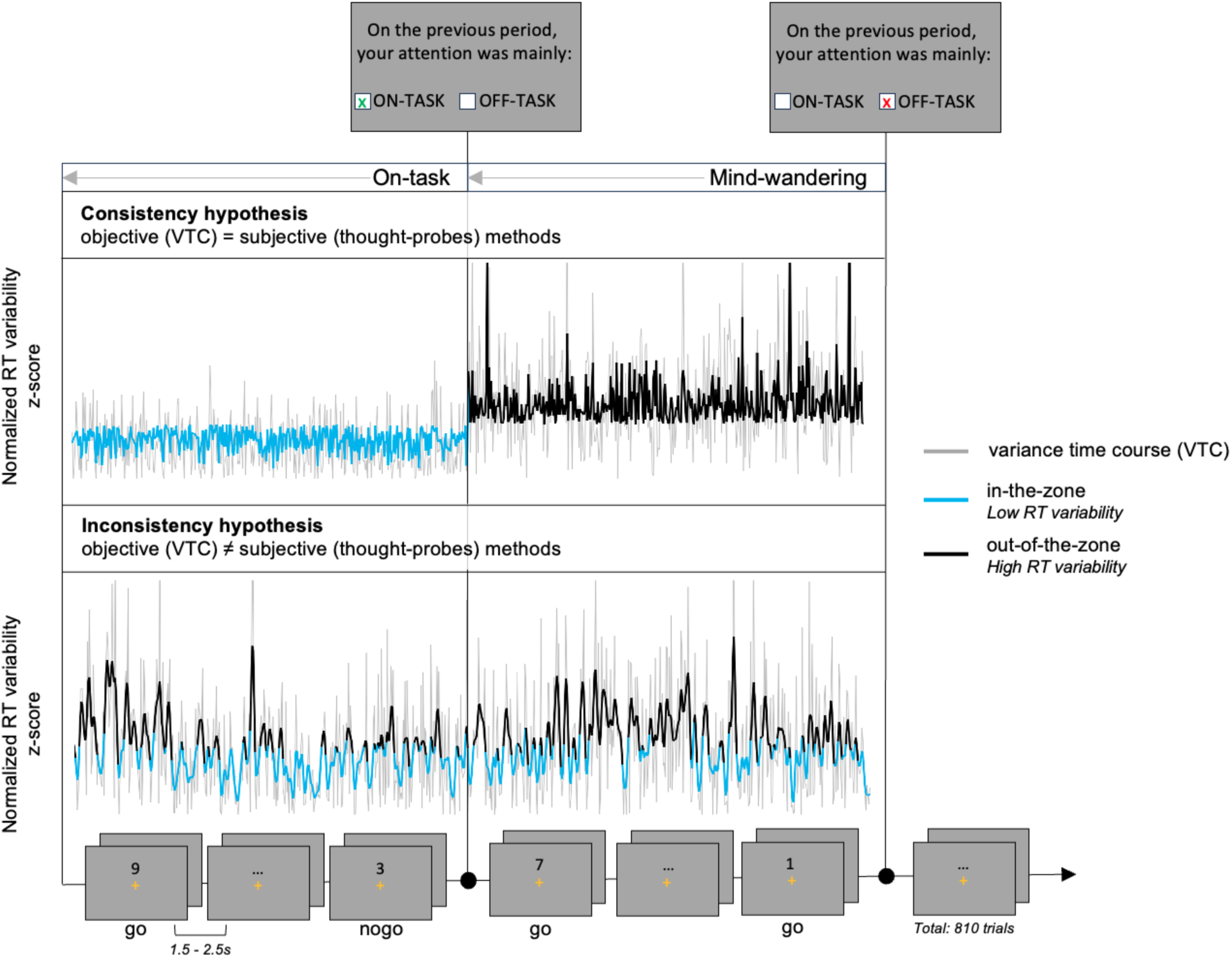
The Sustained Attention to Response Task (SART) and primary hypotheses. Participants are instructed to press every number (go trials) except the number 3 (no-go trials). Thought-probes were used to detect on-task and mind-wandering periods (subjective approach) and the variance time course was used to distinguish periods of low (in-the-zone) and high (out-of-the-zone) RT variability (objective approach). According to the consistency hypothesis, an on-task subjective report should be preceded by a higher time spent in-the-zone whereas a mind-wandering report should be preceded by higher time spent out-of-the-zone. The inconsistency hypothesis predicts no such pattern.

### 2.3. Measuring the fluctuations of sustained attention

#### 2.3.1. The subjective thought-probe methodology

At every 30-trial interval, thought probes were administered to assess whether participants’ attention was primarily on the task (on-task) or diverted to unrelated thoughts (off-task). The SART included 27 such probes in total. The count of no-go trials ranged from 3 to 4 per interval between thought probes (i.e., block). Clear instructions were provided to differentiate between the two states of attention before beginning the task.

#### 2.3.2. The objective Variance Time Course (VTC) methodology

Adopting the approach outlined by Esterman et al. (2013), we calculated intra-individual variability in RTs by determining the variance time course (VTC) for each participant. RTs were first normalized within-subjects to standardize the VTC scale. The VTC was then derived from the accurate go trial responses, where each trial’s value corresponds to the absolute difference between that trial’s RT and the average RT. For error trials and trials lacking responses, such as missed responses on go trials or successfully withheld responses on no-go trials, values were estimated via linear interpolation, based on the RTs from the adjacent trials. The incorporation of thought probes necessitated an adaptation to Esterman et al.’s method, leading us to smooth the VTC within each block rather than across the entire task. This adjustment allowed us to control for potential change in variability of RT induced by the probes. To achieve this within-block smoothing, a Gaussian kernel with a full-width at half-maximum of three trials was used, thereby integrating data from the six nearest trials through a weighted average. Performance was categorized into two segments based on the median of the smoothed VTC: an in-the-zone state with reduced RT variability and an out-of-the-zone state with increased RT variability. Each non-contiguous segment lasted for 15 minutes.

### 2.4. Behavioral analysis

To confirm the effectiveness of both subjective and objective methods in capturing the dynamics of sustained attention, we calculated traditional performance metrics. The average reaction time (mean RT) for correct go trials was measured based on button presses to assess speed. For accuracy, we determined the percentage of attention lapses, operationalized as commission errors. Commission error rate is the ratio of no-go trial errors to the total number of no-go trials within a specified attentional state. The rate of omission errors (failures to press to go trials) was low, at 0.86% and thus these trials were not considered. Our primary analysis centered on three key dependent variables:

#### 2.4.1. The time spent ‘out-of-the-zone’

To quantify the percentage of time participants spent out-of-the-zone, we computed the duration in seconds spent in a state of high RT variability within each subjective state—either on-task or off-task. We then divided this by the total time spent in the corresponding subjective state throughout the entire task. This calculation method allows us to incorporate the number of probes categorized as on-task or off-task into our percentage.

#### 2.4.2. Predictors of attentional lapses

To test whether the two approaches predicted the occurrence of attentional lapses independently and whether each approach explains its own variance, we performed a binomial logistic regression analysis to explore how the subjective and objective states predict the commission errors. We used the variance inflation factor (VIF) to measure the degree of collinearity in our regression model. A VIF below 5 will indicate low collinearity of our predictors and suggests independent variances of the subjective and objective measures (Shrestha, 2020).

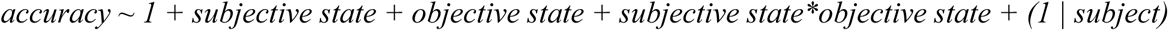

#### 2.4.3. The cognitive control mechanisms: RT around no-go trials

To reproduce the pre-trial slowing analysis of Esterman and colleagues (2013), we analyzed the mean RT of the three correct go trials before a correct inhibition to a no-go trial (the pre-correct trials) and compared them the three go trials before a commission error (the pre-error trials; deBettencourt et al., 2015; deBettencourt et al., 2018). Individuals with no commission errors or no correct inhibition in at least one condition (in-the-zone/on-task; in-the-zone/off-task; out-of-the-zone/on-task; out-of-the-zone/off-task) were excluded from the analysis (*n*=7). Similarly, post-error slowing was explored by comparing mean RTs on the three go trials immediately following a commission error (the post-error trials) to the three go trials that preceded it (the pre-error trials; Dutilh et al., 2012). Rare erroneous go trials were excluded from analysis. If another no-go trial was present in the go periods around the no-go, the trial was excluded from the analysis.

### 2.5. Statistical analysis

The performance measures were subjected to analysis of variance (ANOVA) including the within-subject factors Objective State (in-the-zone vs. out-of-the-zone) and Subjective State (on-task vs. mind-wandering). Within-subject t-tests were used to compare whether the time spent out-of-the-zone was different when the subject declared to be on-task compared to mind-wandering. For the cognitive control engagement, we added to the ANOVA the within-subject factor Period (pre-vs. post-) and Response Type (correct inhibition vs. commission error). Effect sizes were calculated using partial eta square (η2p) for the ANOVA and Cohen’s d for the t-tests. In the case of statistically significant interactions, paired t-tests were used. A two-tailed significance level of 0.05 was used for all tests.

## RESULTS

### 3.1. Overall sustained attention performance

*Commission errors*. The ANOVA performed on the rate of commission errors revealed main effects of both Objective, *F*(1,39) = 130.87, *p* < 0.001, η²p =.77 (**Figure 2A**) and Subjective, *F*(1,39) = 21.32, *p* < 0.001, η²p =.35 states (**Figure 2B**). Error rates were lower during in-the-zone (mean ± sd; 27.06% ± 19.97) compared to out-of-the-zone (45.61% ± 22.10) periods and on-task (31.45% ± 19.47) compared to mind-wandering (41.21% ± 25.18) periods. No significant Objective state x Subjective state interaction was observed (*p* = 0.226).

**Figure 2.**
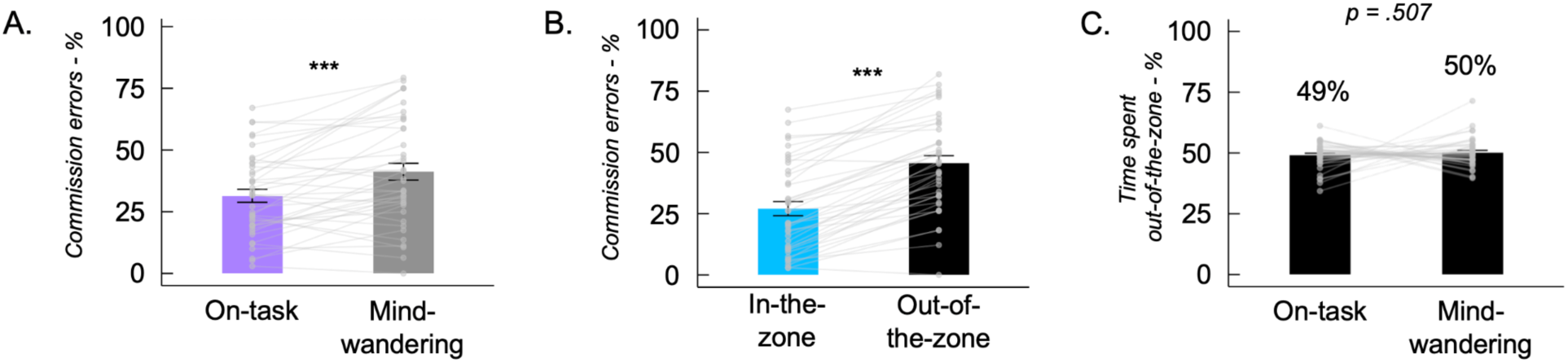
Effect of subjective and objective approaches on performance. A. The subjective approach revealed a higher lapse rate during mind-wandering compared to on-task periods. B. The objective approach revealed a higher lapse rate during out-of-the-zone compared to in-the-zone periods. C. No significant difference in the time spent out-of-the-zone during on-task and mind-wandering periods. Error bars represent standard-errors. ***: p <.001.

*Reaction Times.* The ANOVA performed on mean reaction times revealed a main effect of the objective state, *F*(1,39) = 123.08, *p* < 0.001, η²p =.76, with faster RTs during in-the-zone periods (359 ms ± 7.36) compared to out-of-the-zone periods (381 ms ± 7.77). No main effect of the subjective state (p = 0.891) or objective state x subjective state interaction (p = 0.769) was observed.

### 3.2. Overlap of objective and subjective states

The t-test revealed that the percent of time spent out-of-the-zone was similar between on-task periods (49.0% ± 5.28) and mind-wandering periods (50.1% ± 5.89), *t*(39) =-0.67, *p* = 0.507, d =-0.11, IC 95 % [-0.42, 0.21], suggesting that the subjective and objective approaches did not capture overlapping attention fluctuations (**Figure 2C**). Demonstrating that these results do not rely on temporally smoothing and binarizing VTC values, a t-test performed on the unsmoothed VTC revealed no difference between RT variability during on-task (0.713 ± 0.05) and mind-wandering periods (0.724 ± 0.06), *t*(39) =-0.98, *p* = 0.331, d =-0.16, IC 95 % [-0.47, 0.16].

### 3.3. Predictors of attentional lapses

The occurrence of attentional lapses was predicted by both subjective state, *b* =-.44, SE=.10, *p* < 0.001; and objective state, *b* =.91, *SE*=.096, *p* < 0.001. However, no subjective x objective state interaction was observed, *b* = 0.05, *SE* =.192, *p* = 0.794. Interestingly, the variance inflation factor (VIF) that measures the degree of collinearity in our regression model was below 5 for the subjective states (VIF = 2.13), the objective states (VIF = 2.02), and the subjective x objective states interaction (VIF = 3.12), suggesting that the variables do not correlate with each other and explain unique variance in attentional lapse rates.

### 3.4. RTs dynamics around no-go trials

Another way to explore whether the objective and subjective approaches reflect common attention fluctuations is to compare their effects on RTs before and after no-go trials. The ANOVA performed on mean RTs revealed main effects of Period, *F*(1,32) = 7.03, *p* = 0.012, η²p =.18, Response Type, *F*(1,32) = 37.51, *p* < 0.001, η²p =.54, and Objective state, *F*(1,32) = 24.82, *p* < 0.001, η²p =.44, but no main effect of Subjective state, *F*(1,32) = 1.93, *p* = 0.175. RTs were slower after than before no-go trials and were slower around correct inhibitions compared to commission errors. RTs were also slower during out-of-the-zone compared to in-the-zone periods (see supplementary results for intermediate interactions). Interestingly, the ANOVA revealed an interaction between the four independent variables, suggesting the objective and subjectives states interact on the RTs around no-go trials, *F*(1,32) = 7.03, *p* = 0.012, η²p =.18. To describe this interaction, we characterized pre-error speeding and post-error slowing with post-hoc t-tests.

#### 3.4.1 Precursors of correct inhibition versus commissions errors

Previous studies have revealed slower RTs before correct inhibitions compared to commission errors (deBettencourt et al., 2015; deBettencourt et al., 2018). In the current study, the post-hocs tests replicated these results with slower RTs on the trials preceding a correct inhibition compared to a commission error but only during out-of-the-zone periods: out-of-the-zone and on-task, *t(*32) =-4.95, *p* < 0.001, *p_tukey_*= 0.002 (**Figure 3A**); out-of-the-zone and mind-wandering, *t(*32) =-4.18, *p* < 0.001, *p_tukey_*=0.015, (**Figure 3B**). The effect did not survive to correction for multiple comparisons during in-the-zone periods: in-the-zone and on-task, *t(*32) =-2.79, *p* = 0.009, *p_tukey_*= 0.323; in-the-zone and mind-wandering, *t(*32) =-3.46, *p* = 0.002, *p_tukey_*= 0.088.

**Figure 3.**
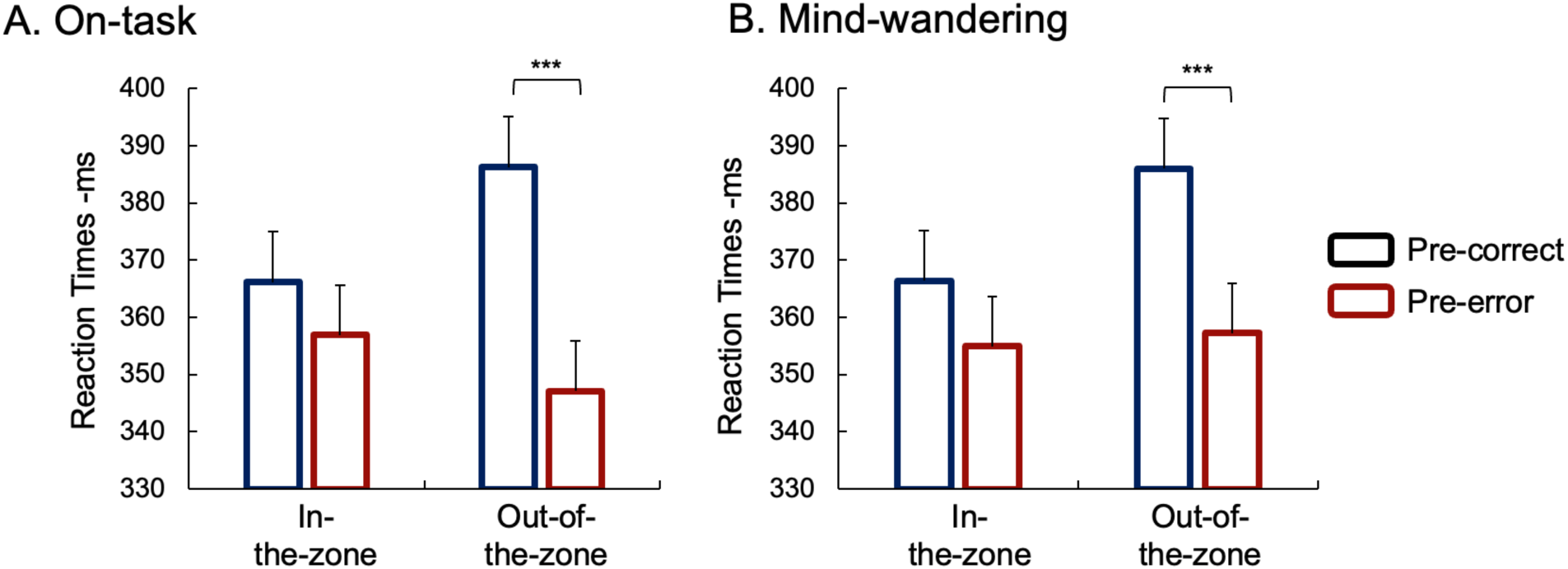
Results on the mechanism of attentional preparation. The objective method revealed that RTs were slower before correct vs. incorrect inhibitions during periods of high RT variability, suggesting higher anticipation of no-go trials during out-of-the-zone states. No effect of the subjective on-task (A) and mind-wandering periods (B) have been observed on attentional preparation. Error bars represent standard-errors. ***: p <.001.

Interestingly, the slowing of RTs preceding a correct inhibition was more pronounced in the out-of-the-zone state than in the in-the-zone state, both when subjects reported being on-task, *t*(32) =-3.43, *p* = 0.002, and mind-wandering, *t*(32) =-3.64, *p* < 0.001. However, these effects did not survive multiple correction procedures (on-task, *p_tukey_*= 0.094; mind-wandering, *p_tukey_*= 0.058). No other effect has been observed (all p’s > 0.05).

#### 3.4.2 Post-error slowing

When participants reported being subjectively on-task and were objectively out-of-the-zone, RTs were slower after than before errors, indicating post-error slowing, *t(*32) =-3.90, *p* < 0.001, *p_tukey_*= 0.032. However, no post-error slowing was observed during the three other combined attentional states (all *p* > 0.1), suggesting that the objective and subjective states interact to modulate the conflict adaptation response (**Figure 4A, 4B**).

**Figure 4.**
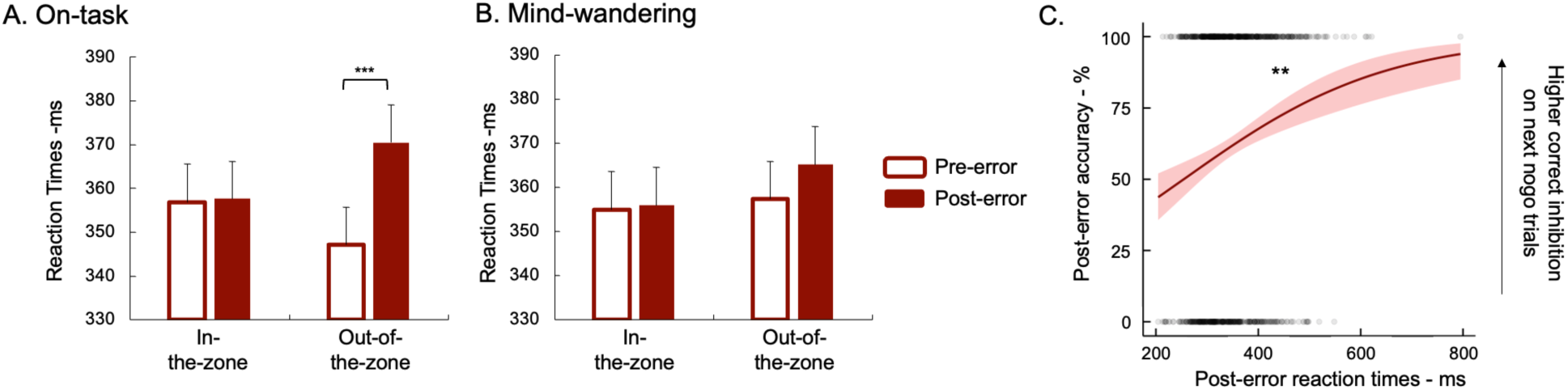
Results on the adaptation control mechanism: post-error slowing. Post-error slowing was only observed during the combined out-of-the-zone/on-task period (A). No post-error slowing was observed during mind-wandering (B). Longer RTs after a commission error was associated with higher accuracy on the next no-go trials (C). Error bars represent standard-errors. **: p <.01; ***: p <.001.

Surprisingly, when participants reported being subjectively in an mind-wandering state and were objectively out-of-the-zone, RTs were numerically slower after than before correct inhibitions, indicating a post-correct slowing, *t(*32) =-2.09, *p* = 0.045, although this effect did not survive to the correction for multiple comparisons (*p_tukey_*= 0.757). No post-correct slowing was observed during the three other combined attentional states (all *p* > 0.1).

To further explore the role of post-error slowing in performance, we conducted a binomial logistic regression analysis to investigate how the reaction times following a commission error predict the accuracy of the subsequent no-go trial, while controlling for the number of trials between two no-go presentations. The model was specified as follows:

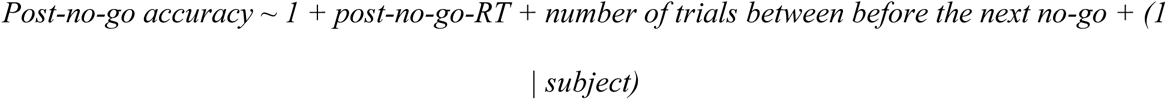

The analysis revealed that greater post-error slowing predicts better correct inhibition on the next no-go trial, *b* =-.00461, *SE* =.0014, *p* = 0.001; with no effect of the number of trials before the next no-go trial, *b* =-.00261, *SE* =.0215, *p* = 0.904 (**Figure 4C**). This result reveals a key role of post-error slowing to restore attentional engagement on the task and avoid future errors.

## DISCUSSION

The goal of this study was to investigate the degree of consistency between objective and subjective measures of sustained attention fluctuations. For the first time, we integrated approaches tracking attention fluctuations with the VTC and mind-wandering probes within a single study. Replicating existing literature, participants made more frequent attentional lapses during out-of-the-zone periods characterized by high response time variability and off-task periods associated with mind-wandering. Participants, however, were no more likely to be out-of-the zone when mind wandering vs. on task, and objective and subjective states explained unique variance in lapse rates. Thus, our findings demonstrate that objective and subjective measures of sustained attention do not fluctuate concurrently and provide evidence that these approaches may capture distinct types of attention fluctuations.

In this study, the hypothesis of inconsistency between objectively and subjectively measured fluctuations was initially supported by the analysis of overlap between the two. The percentage of time spent out-of-the-zone was similar in both on-task (49.0%) and mind-wandering states (50.1%). This initial result suggests that periods of poor objective and subjective attention do not occur simultaneously and seems to contradict the recent findings of Yamashita et al. (2021). Based on fMRI data, the authors identified brain states whose behavioral patterns are identical to in-the-zone and out-of-the-zone behavioral periods. Collecting self-reports of mind-wandering on a scale from 0 (only on-task) to 100 (only off-task), they found that participants reported being off-task more when in-the-zone than out-of-the-zone (Yamashita et al., 2021). This supports an “inverse consistency” hypothesis where the in-the-zone state reflects a mind-wandering state. However, if participants primarily reported mind-wandering when in-the-zone, one would expect scores above 50 on the mind-wandering scale. Yet, participants’ average ratings were around 30 on the mind-wandering scale when in-the-zone, suggesting they primarily reported being on-task rather than mind wandering. Reporting being less on-task does not necessarily suggest off-task thought or mind wandering, but could reflect an instability in actively maintaining the task set in mind. It is therefore possible that the in-the-zone state is characterized by a less effortful, costly, or conscious activation of the task-set compared to out-of-the-zone periods requiring more control, with both states being independent from the occurrence of mind-wandering episodes.

A second piece of evidence supporting the inconsistency hypothesis relates to the predictors of attentional lapses. Our logistic regression model highlighted that the objective and subjective measures were not collinear and instead independently predicted attentional lapses. This within-subject result complements those of Unsworth and colleagues (2021) at the between-subject level. Their between-subject latent variable analysis revealed that reaction time variability on the SART, operationalized by the coefficient of variation, and self-reported task-unrelated thoughts represent two independent and distinct measures.

A third piece of evidence supporting the hypothesis of inconsistency comes from the engagement of cognitive control mechanisms as indicated by attentional preparation observed before a correct no-go response. Our results replicate prior observations of longer RTs before a correct inhibition compared to an error during periods of high but not low RT variability (Esterman et al., 2013). This suggests that cognitive control is more engaged during out-of-the-zone than in-the-zone periods. If the consistency hypothesis were true we would observe greater attentional preparation before a correct response only during subjective mind-wandering periods and not during on-task periods. However, our post-hoc analysis, as well as the Period x Response Type x Subjective States interaction (in the supplementary section), shows that attentional preparation with a correct response is observed in both subjective states and not just during mind-wandering periods, suggesting that the preparation mechanisms are not modulated by the subject’s subjective state but only by the objective attentional state based on RT variability.

The results on adaptive control measured with post-error slowing also support the inconsistency hypothesis and provide initial evidence for a “combined states” hypothesis that objectively and subjectively measured fluctuations interact to predict control. Indeed, our results reveal an interaction between objective and subjective states and show that post-error slowing is only observed during simultaneous on-task/out-of-the-zone periods, but not in the other three combined states. The presence of post-error slowing only in the on-task/out-of-the-zone state could reflect the disengagement of cognitive control before an error in this state. This is consistent with previous fMRI results showing a reduction in dorsal attention network activity before an error during out-of-the-zone periods (Esterman et al., 2013). The slowing after an error would thus reflect the re-engagement of control preparation for the next no-go trial. This performance restoration is evidenced by our logistic regression analysis, which shows that long RTs after an error predict correct inhibition on the next no-go trial. The absence of post-error slowing in the on-task/in-the-zone state could suggest preserved cognitive control throughout this period with commission errors arising through cognitive mechanisms other than control failures.

If errors during in-the-zone periods are not characterized by a reduction in cognitive control, it is therefore coherent that post-error slowing is not observed regardless of whether the subject reports being on or off task. Why, however, is no post-error slowing observed in the out-of-the-zone state when the subject is mind-wandering? One potential explanation could be error awareness. Previous studies have shown that post-error slowing is only observed when subjects are aware of their errors (Klein et al., 2007; Nieuwenhuis et al., 2001). During off-task periods, it is therefore possible that the participant, occupied by mind-wandering, does not become aware of the error, thus preventing the re-engagement of control over the ongoing task.

Overall, these results support the inconsistency hypothesis that self-reported mind wandering and performance fluctuations measured with RT variability capture different sustained attentional dynamics. This may explain the contradictory findings in the literature, particularly the observation that default mode network activity is higher during both mind-wandering *and* in-the-zone periods of performance. That is, periods of error-prone mind-wandering have been associated with increased DMN activity, whereas the objective approach links the DMN to greater behavioral stability and fewer errors. This contrast makes it challenging to directly link DMN activity with task performance and to understand its role in sustained attention. The inconsistency hypothesis, supported by the results of the current study, reconciles these findings by suggesting that the two approaches measure different types of attentional fluctuations and might explain why different patterns of DMN activity were observed in the literature, with some results associating DMN with poor performance and others with good performance. This interpretation is also supported by the findings of Kucyi et al. (2016), who attempted to disentangle the variance underlying DMN activity. They revealed that both mind-wandering thoughts and behavioral response time stability are both related to DMN activity, suggesting that the DMN supports multiple processes.

## LIMITATIONS

Although these different results converge and suggest that RT variability and mind-wandering reflect different phenomena in CPTs and inhibition tasks, it is possible that, in some tasks, RT variability may be more strongly correlated with periods of mind-wandering. For instance, an increasing number of studies show that in the Metronome Response Task, RT variability reliably and validly predicts mind-wandering (Anderson et al., 2021; Seli et al., 2013). In another recent study, Jalava and Wammes found that RT variability and speed predict depth of mind-wandering (Jalava & Wammes, 2024). However, this task remains limited as it does not allow for the demonstration of impaired attentional performance during periods of mind-wandering and the increased occurrence of attentional lapses. Furthermore, since this task requires only a single response, RT variability may not reflect the competition that might exist between prepotent responses observed in go/no-go tasks, but rather direct fluctuations in task-related thoughts. The existence of multiple sources of RT variability will need to be taken into account in future studies.

## CONCLUSION

To conclude, all of our results, from analyzing the time spent out-of-the-zone during mind-wandering periods to predictors of attentional lapses to control mechanisms, unanimously support that the objective and subjective approaches do not measure the same types of fluctuations in sustained attention during go/no-go tasks. Future studies should attempt to better characterize the nature of these two forms of attentional fluctuations and delineate the different cognitive processes that underlie them. Overall, these results underscore the importance of integrating both objective and subjective measures to fully characterize the rich dynamics of attentional states fluctuations. Solely relying on one approach may only capture part of the picture, potentially leading to contradictory conclusions across studies. Only by understanding how objective cognitive performance and subjective experience diverge and interact can we fully elucidate the complex attentional processes allowing us to remain focused on goals despite internal and external distraction.

## DATA AVAILABILITY STATEMENT

The data are available on the Open Science Framework repository (https://osf.io/xwe23/).

## ACKNOWLEDGEMENTS

This research was supported by a French Ministry of Higher Education and Research grant that was allocated to the first author and Office of Naval Research grant N00014-23-1-2768 to E.K.V. and M.D.R. We thank the staff at Unit 1114 of the French National Institute for Health and Medical Research and at the University of Strasbourg.

## AUTHOR CONTRIBUTION

**Matthieu Chidharom**: Conceptualization, Data Curation, Formal Analysis, Methodology, Project Administration, Supervision, Validation, Visualization, Writing-Original Draft, Writing-review & Editing. **Anne Bonnefond**: Resources, Writing-review & Editing. **Edward K. Vogel**: Resources, Supervision, Writing-review & Editing. **Monica D. Rosenberg**: Resources, Supervision, Writing-review & Editing.

